# Amyloid-beta pathology increases synaptic engulfment by glia in feline cognitive dysfunction syndrome: A naturally occurring model of Alzheimer’s disease

**DOI:** 10.1101/2025.03.06.641819

**Authors:** Robert I. McGeachan, Lucy Ewbank, Meg Watt, Lorena Sordo, Alexandra Malbon, Muhammad Khalid F. Salamat, Makis Tzioras, Joao Miguel De Frias, Jane Tulloch, Fiona Houston, Danièlle Gunn-Moore, Tara Spires-Jones

## Abstract

Feline cognitive dysfunction syndrome (CDS) is an age-related neurodegenerative disorder, comparable to dementia in people, characterised by behavioural changes such as increased vocalisation, altered social interactions, sleep-wake cycle, disorientation and house-soiling. Although the underlying mechanisms remain poorly understood, pathologies similar to those observed in Alzheimer’s disease (AD), have been identified in the brains of aged or CDS-affected cats, including brain atrophy, neuronal loss, amyloid-beta plaques, tau pathology, and cerebral amyloid angiopathy. Neuroinflammation and synapse loss, other important hallmarks of AD, may also play important roles in feline ageing and CDS, but these are yet to be explored. Several mechanisms of synapse loss have been described in human AD and mouse models of amyloidopathy, including synaptic accumulation of amyloid-beta, and the aberrant induction of synaptic engulfment by microglia and astrocytes. In this study, immunohistochemistry and confocal microscopy were used to examine the parietal cortex of young (n=7), aged (n=10), and CDS-affected (n=8) cats. Linear mixed effect modelling revealed that amyloid-beta accumulates within synapses in the aged and CDS-affected brain. Additionally, in the aged and CDS groups there was microgliosis, astrogliosis and increased synaptic engulfment by microglia and astrocytes in regions with Aβ plaques. Further, microglia and astrocytes show increased internalisation of amyloid-beta-containing synapses near plaques. These findings suggest that amyloid-beta exerts a pathogenic effect in the feline brain, with mechanisms mirroring those seen in human AD.

## Introduction

Feline cognitive dysfunction syndrome (CDS) is a clinical disorder observed in elderly cats which is similar to human dementia. CDS is characterised by behavioural changes, including increased vocalisation, altered social interactions, changes in the sleep-wake cycle, disorientation, and house-soiling. In practice, feline CDS is likely underreported and often unrecognised or underdiagnosed by both owners and primary care veterinary surgeons. There are limited studies investigating the prevalence of feline CDS, but one survey found that 28% of cats aged 11–14 years exhibited at least one clinical sign of CDS, with this increasing to 50% of cats aged over 15 years (Moffat & Landsberg, 2003).

The pathophysiology of feline CDS is poorly understood. However, pathologies comparable to those observed in human Alzheimer’s disease (AD), the most common cause of human dementia, have been identified in the aged or CDS- affected feline brain, including brain atrophy and neuronal loss (Levine *et al*., 1986; Levine, 1988; Zhang *et al*., 2005, 2006; Chambers *et al*., 2015), amyloid-beta plaques (Cummings *et al*., 1996; Nakamura *et al*., 1996; Brellou *et al*., 2005; Head *et al*., 2005; Gunn-Moore *et al*., 2006, 2007; Sordo *et al*., 2021), tau pathology (Head *et al*., 2005; Gunn-Moore *et al*., 2006, 2007; Chambers *et al*., 2015), and vascular disease including cerebral amyloid angiopathy (Cummings *et al*., 1996; Nakamura *et al*., 1996; Head *et al*., 2005; Gunn-Moore *et al*., 2007). The amyloid-beta deposits in cats are often less mature and more diffuse than the dense-core plaques observed in human AD. Instead, these diffuse plaques more closely resemble the amyloid-beta pathology found in the brains of healthy, aged humans (Gunn-Moore *et al*., 2007; Zaletel *et al*., 2021). Additionally, similar to findings in human AD (Morris *et al*., 2014), the degree of amyloid pathology correlates poorly with behavioural and cognitive changes in patients with CDS (Head *et al*., 2005). Furthermore, although studies have shown an age-related increase in amyloid-beta accumulation in the feline brain, one study found no significant increase in amyloid-beta pathology in cats with CDS compared to age-matched controls (Sordo *et al*., 2021). This raises an important question: is amyloid-beta pathology simply an age-related incidental finding, or does it actively contribute to neurodegeneration and the signs observed in feline CDS? Determining whether amyloid-beta exerts a pathogenic effect in the feline brain is becoming increasingly relevant, particularly as monoclonal antibodies targeting amyloid-beta, which have shown modest success in slowing cognitive decline in people, are being approved for the treatment of AD across the world (Sims *et al*., 2023; van Dyck *et al*., 2023)

In human AD, there are pronounced inflammatory changes in microglia and astrocytes, including changes in gene expression, morphology, and increased density of these cells, with the most severe changes in the direct vicinity of amyloid plaques (Keren-Shaul *et al*., 2017; Henstridge, Hyman, *et al*., 2019; Habib *et al*., 2020). While gliosis was initially thought to be largely downstream of pathological changes in neurons, substantial genetic data indicate changes in genes expressed in astrocytes and microglia are associated with increased risk of AD. Further, recent data link both microglia and astrocytes to synapse loss (Hong *et al*., 2016a; Tzioras, Daniels, *et al*., 2023; Tzioras, McGeachan, *et al*., 2023), which is the strongest pathological correlate of cognitive decline (DeKosky & Scheff, 1990; Terry *et al*., 1991; Scheff & Price, 2006; Scheff *et al*., 2007). Understanding and targeting the mechanisms driving glial changes and their contributions to synapse loss are therefore considered a promising therapeutic approach for slowing or halting the progression of AD. Several mechanisms of synapse loss have been described (Tzioras, McGeachan, *et al*., 2023).

One proposed mechanism is that amyloid-beta is directly synaptotoxic. Post-mortem analyses of AD brains (Koffie *et al*., 2012; Jackson *et al*., 2019), along with studies using preclinical mouse models with amyloid plaque deposition (Koffie *et al*., 2009; Pickett *et al*., 2019), suggest that oligomeric amyloid-beta accumulates directly within synapses. This accumulation is particularly prominent near extracellular amyloid plaques, where significant excitatory synapse loss is also observed (Koffie *et al*., 2009, 2012). Another potential mechanism leading from pathological amyloid-beta accumulation to synapse loss is via aberrant induction of synaptic pruning by glia. Synaptic pruning mediated by astrocytes and microglia is an important aspect of normal circuit development (Schafer *et al*., 2012; Chung *et al*., 2013; Cserép *et al*., 2020; Lee *et al*., 2021). Recently, astrocytes and microglia were observed to engulf synapses in post-mortem AD brain, with the greatest levels of engulfment in the vicinity of amyloid plaques (Tzioras, Daniels, *et al*., 2023). Increased synaptic engulfment by glia isn’t specific to Alzheimer’s disease and has been described in schizophrenia other neurodegenerative disease (Henstridge, Tzioras, *et al*., 2019; Tzioras *et al*., 2021; Dando *et al*., 2024; McGeachan *et al*., 2024). Although it is now well established that amyloid-beta pathology accumulates in the aged and CDS- affected feline brain (Sordo *et al*., 2021), the relationship to synapse degeneration has not been explored.

This study investigates whether amyloid-beta pathology in the feline brain accumulates within synapses and is associated with increased synaptic engulfment by glia.

## Methods

### Post-mortem samples and ethical approval

The cat brains used in this study were obtained from The Royal (Dick) School of Veterinary Studies. Cats were diagnosed as having CDS if they exhibited behavioural changes consistent with CDS for at least three months that could not attributed to any other medical condition (Gunn-Moore *et al*., 2007). CDS-affected cats (mean age = 16.9 years) were compared to “aged” (mean age = 17.17 years) and “young” (mean age = 4.81 years) control groups with no history of neurologic disease. The CDS and “aged” group were age-matched (Wilcoxon rank sum test, W=36.5, *p* = 0.788). Two-thirds of the cats were female (64%; n=16), one-third male (32%, n=8), and one cat was of unknown sex. The CDS, age-matched and young groups were sex-matched (Fisher’s exact test, *p* = 0.34). The demographic details of all cases used in this study, stratified by group, are detailed in **Table 1**. Consent was obtained from all owners and rescue shelters for the donation of the feline’s bodies for research purposes. Ethical approval was gained via the Veterinary Ethical Review Committee from the RDSVS, The University of Edinburgh (VERC 50.17 and 30.20). The brains were fixed in 10% buffered formalin, dissected into anatomical brain regions and then embedded into paraffin blocks before being cut to a thickness of 4μm and mounted onto glass slides. The focus was to examine the parietal cortex as a previous study has demonstrated the presence of amyloid-beta within this brain region of aged and CDS feline brains (Sordo *et al*., 2021).

**Table 1.** Case details. CDS = cognitive dysfunction syndrome. M = Male, F = Female,= sex unknown

| Young (N=7) |  |  | Aged without CDS (N=10) |  |  | CDS (N=8) |  |  |
| --- | --- | --- | --- | --- | --- | --- | --- | --- |
| ID | Age | Sex | ID | Age | Sex | ID | Age | Sex |
| Cat 1 | 2 | F | Cat 8 | 14 | M | Cat 18 | 11 | F |
| Cat 2 | 4 | F | Cat 9 | 14 | M | Cat 19 | 14 | F |
| Cat 3 | 4 | F | Cat 10 | 15 | F | Cat 20 | 16 | F |
| Cat 4 | 5 | ? | Cat 11 | 15 | F | Cat 21 | 18 | F |
| Cat 5 | 6 | F | Cat 12 | 16 | F | Cat 22 | 19 | M |
| Cat 6 | 6 | M | Cat 13 | 16 | M | Cat 23 | 19 | M |
| Cat 7 | 7 | F | Cat 14 | 17 | F | Cat 24 | 19 | F |
|  |  |  | Cat 15 | 19 | F | Cat 25 | 20 | F |
|  |  |  | Cat 16 | 20 | M |  |  |  |
|  |  |  | Cat 17 | 26 | M |  |  |  |

### Immunohistochemistry

The paraffin-embedded tissue slides were dewaxed in xylene for 6 minutes and rehydrated in ethanol-to-water solutions with decreasing concentrations (100%, 90%, 70%, 50%) for 3 minutes each to rehydrate the tissue, then thoroughly rinsed with 100% water. Antigen retrieval was performed by immersing slides in 90% formic acid for 5 minutes at room temperature followed by citrate buffer (Vector labs, H3300) for 3 minutes in a pressure cooker set to the steam setting. Slides were rinsed in water for 5 minutes. To reduce autofluorescence (such as that originating from lipofuscin) the autofluorescence eliminator reagent (Merck, 2160) was used as per the manufacturer’s instructions. The tissues were outlined using a wax pen, followed by washing with PBS for 5 minutes, then washed with PBS and 0.3% Triton-X for an additional 5 minutes. To eliminate autofluorescence originating from red blood cells the Vector TrueView Autofluorescence Quenching Kit (SP-8400-15) was used as per the manufacturer’s instructions. After a 5-minute wash in PBS, the sections were incubated in blocking solutions consisting of PBS, 0.3% Triton X-100, and 10% normal donkey serum for 1 hour. Subsequently, the sections were incubated overnight at 4°C with the primary antibodies (**Table 2**). The following day, the slides underwent two 5-minute washes using PBS with 0.3% Triton X-100, a 5-minute wash with PBS, and were then incubated with the secondary antibodies (**Table 2**) for 1 hour at room temperature. To remove unbound secondary antibodies, the slides were washed twice with PBS and 0.3% Triton X-100, followed by a 5-minute wash with PBS. Finally, one drop of Immumount (Epredia, 9990402) was used in the mounting of the slides to glass coverslips. For each stain, a ‘no primary’ antibody negative control was included.

**Table 2.** Antibody information.

|  | GFAP | 4G8 | SYN1 | IBA1 |
| --- | --- | --- | --- | --- |
| **Primary Antibodies** |  |  |  |  |
| **Species** | Chicken | Mouse (IgG2b) | Rabbit | Goat |
| **Mono/polyclonal** | Polyclonal | Monoclonal | Polyclonal | Polyclonal |
| **Company** | Abcam | Biolegend | Merck | Abcam |
| **Catalogue / lot Number** | cat: #ab4674,<br>lot: GR345545 | cat: #800703,<br>lot: B273175 | cat: #ab1543P,<br>lot: 3993513 | cat: #AB5076,<br>lot: 1053204-1 |
| **Dilution** | 1:1000 | 1:1000 | 1:500 | 1:500 |
| **Secondary Antibodies** |  |  |  |  |
| **Species** | Donkey anti-chicken 405 | Donkey anti-mouse IgG 488 | Donkey anti-rabbit 594 | Donkey anti-goat 647 |
| **Catalogue Number** | cat: #703-475-155,<br>lot: 155983 | cat: #ab150109,<br>lot: GR3228235 | cat: #a21207,<br>lot: 2747441 | cat: #ab150135,<br>lot: GR3286128-1 |
| **Dilution** | 1:500 | 1:500 | 1:500 | 1:500 |
| **Table 2 Antibody information** |  |  |  |  |

### Confocal microscopy and quantitative image analysis

Images were obtained using a 63x oil immersion objective on a confocal microscope (Leica SP8). The imaging parameters were kept consistent throughout each experiment. The tissues were first scanned for amyloid-beta plaques, and where present, an image stack was acquired at the site of pathology and directly adjacent in a region that did not contain amyloid-beta pathology but was still within the same cortical layer and only 1 field of view across from the pathology. In cases without amyloid-beta pathology, 10 image stacks were randomly obtained across the 6 cortical layers of the grey matter. An image was taken of the no primary negative control for each staining batch to ensure there was not non-specific staining. Using custom Image J and MATLAB scripts, the image stacks were segmented, and the percentage volume of the image stack occupied by each channel and the colocalisation between channels was calculated. All custom software scripts used in the image processing and analysis are available on GitHub https://github.com/Spires-Jones-Lab. The experimenter was blinded to the case information during immunostaining, image acquisition and image processing.

### Statistical analysis

Statistical analysis was performed in R studio with R version 4.3.2. The majority of statistical analyses used linear mixed effects models (LMEM)(‘lme4’ R package), as this allowed the testing of whether the group (young vs aged vs CDS) or the presence of amyloid-beta pathology (at plaque vs no plaque) impacted the variable of interest while controlling for potentially confounding variables, such as age, and including random effects to prevent pseudoreplication. The decision to ultimately include potentially confounding fixed and random effects in the model, and prevent overfitting of the model, was done by inspecting Akaike Information Criterion (AIC) and Bayesian Information Criterion (BIC) to assess which model best fits the data. Linear mixed effect models assume linearity, normal distribution of residuals and homogeneity of variance. Linearity was assessed by plotting model residuals against predictors, normality of residuals was checked with a QQ-plot, and homogeneity of variance checked by plotting residuals against fitted values. If the model did not meet the assumptions, data was transformed using the method that transformed each individual model to fit the assumptions. Data transformations tested included square root, log, arcsine square root, and Tukey transformation. Post-hoc testing was conducted for pairwise comparisons (’emmeans’ package), with p-values adjusted using Tukey correction for multiple comparisons. Degrees of freedom were calculated using Kenwood-Roger approximation. Statistics are reported in the results text for main figures and in figure legends for supplementary data.

## Results

### Amyloid-beta colocalises with synapses in the aged and CDS feline brain

Synaptic accumulation of amyloid-beta and the engulfment of synapses by glial cells were assessed by immunostaining for amyloid-beta (4G8), synapses (synapsin-1), astrocytes (GFAP), and microglia (IBA1) (**Figure 1a, 2a, 3a),** along with the quantification of the percentage volume of the 3D image stacks occupied by each individual channel and the colocalisation between channels. First, we set out to investigate if amyloid-beta accumulates within synapses in the aged and CDS cat brain. Linear mixed effect modelling of Tukey transformed data (Variable ∼ Group + 1|Case) revealed that the amount of amyloid-beta (4G8) staining was greater in the CDS (*t*(22) = 4.238, *p* < 0.001) and aged group (*t*(22) = 2.847, *p* = 0.024) when compared to the young group (**Figure 1b**). There was no difference between the CDS and aged group (*t*(22) = 1.667, *p* = 0.240). Similarly, there was increased colocalisation between synapses (synapsin-1) and amyloid-beta (4G8) in both the CDS (*t*(22) = 5.004, *p* < 0.001) and aged groups (*t*(22) = 3.701, *p* = 0.003) compared to the young controls, but no difference between CDS and aged group (*t*(22) = 1.616, *p* = 0.26)((**Figure 1c**).

**Figure 1:**
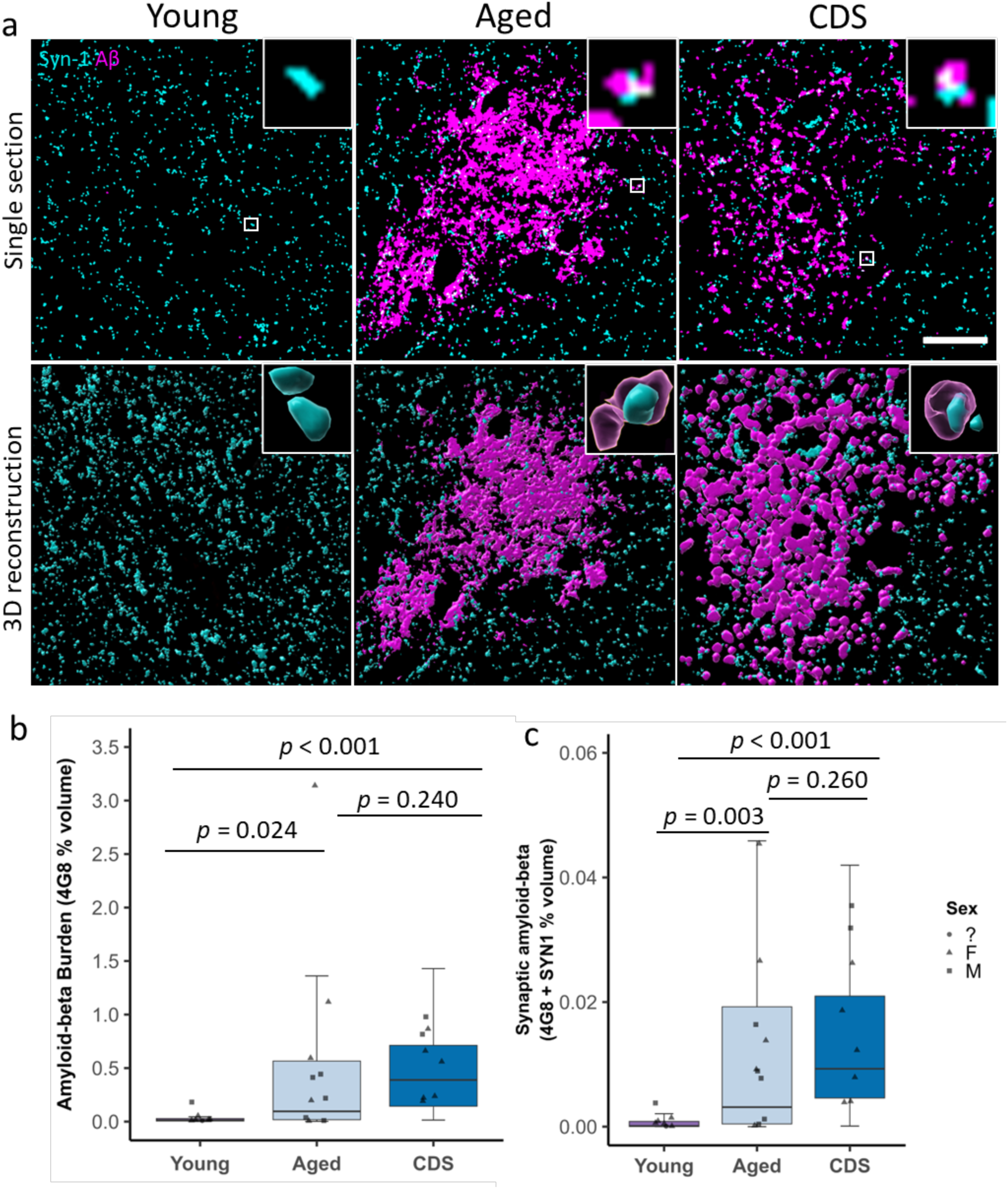
Amyloid-beta colocalises with synapses in the aged and CDS- affected feline parietal cortex. **a)** Segmented confocal images of young, aged, and CDS-affected feline parietal cortex immunostained for synapses (synapsin-1, cyan) and amyloid-beta (4G8, magenta). Scale bar = 20 µm. Inserts = 2 µm * 2 µm. The bottom row of images shows 3D reconstructions generated using IMARIS and have been rotated along the z-axis. Inserts are close up images of the colocalisation highlighted by the white box. **b-c)** Quantitative analysis reveals an increase in amyloid-beta burden (b) and colocalisation between amyloid-beta and synapses (c) in the aged and CDS groups when compared to young controls. Boxplots show quartiles and medians calculated from each image stack. Data points represent case means (sex unknown = circles, females = triangles, males = squares). Statistics: LMEM (variable ∼ Group + 1 | case). *p*-values were calculated from post-hoc testing with Tukey correction for multiple comparisons.

**Figure 2:**
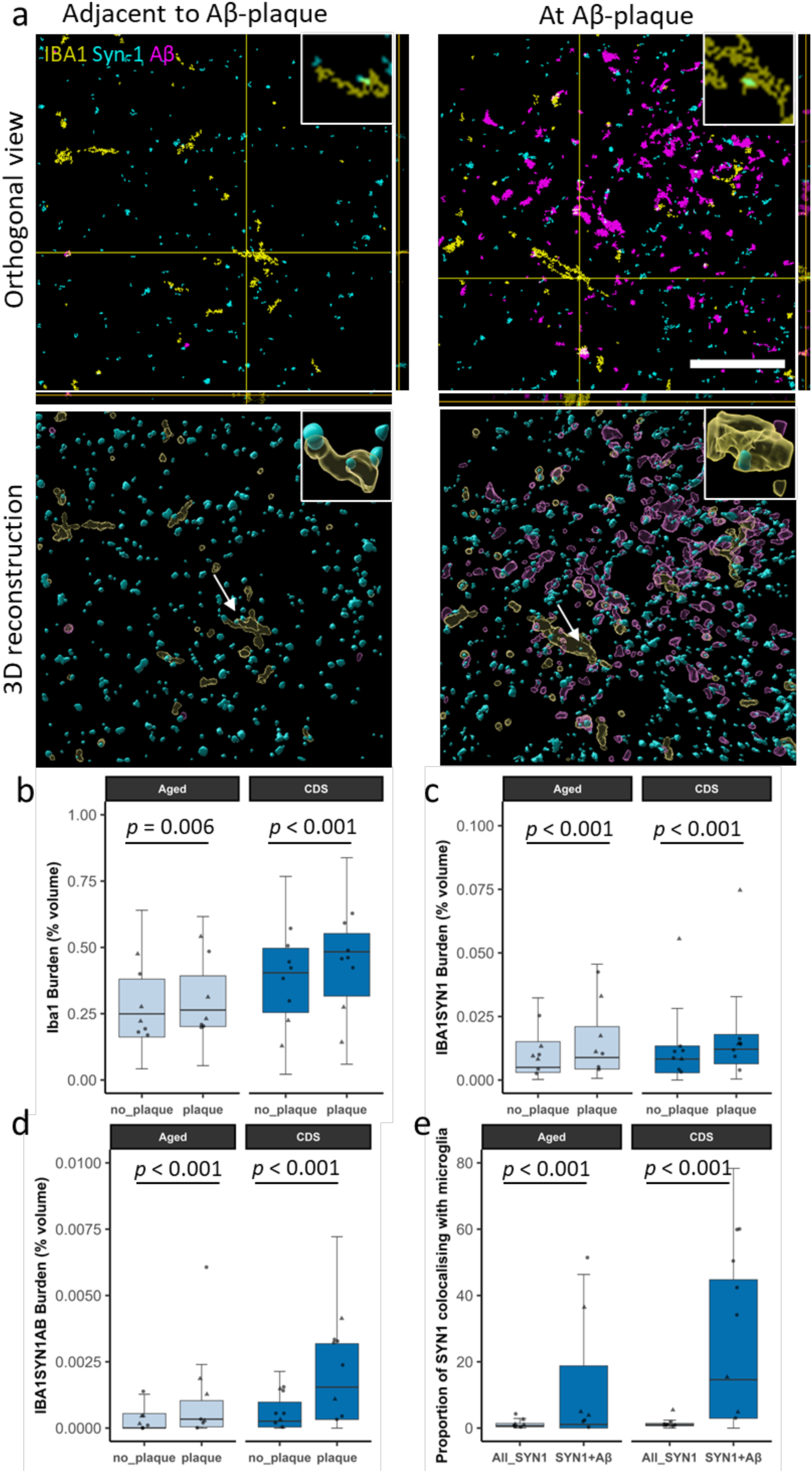
Amyloid-beta pathology induces increased synaptic engulfment by microglia in the feline brain. **a)** Segmented confocal images of CDS-affected feline parietal cortex, showing a region with amyloid-beta pathology and a paired image acquired approximately one field of view away in the same cortical layer in an area without pathology. Tissue was immunostained for synapses (synapsin-1, cyan), amyloid-beta (magenta), and microglia (IBA1, yellow). Scale bar = 20 µm. Insets = 5 µm × 5 µm. The top row of images shows orthogonal views. The bottom row of images shows 3D reconstructions generated using IMARIS and have been rotated along the z-axis, to further demonstrate that synaptic protein is within microglia. Inserts show close up of colocalisation highlighted by arrow. **b-e)** Colocalisation analysis reveals that the image stacks acquired in regions with amyloid-beta pathology show increased microgliosis (IBA1 burden) (b), colocalisation between microglia and synapses (IBA1 and Synapsin-1 burden) (c), and triple colocalisation between microglia, synapses and amyloid-beta (IBA1, synapsin-1 and amyloid-beta burden) (d). Finally, when analysing all image stacks, synapses colocalising with amyloid-beta are more likely to be engulfed by microglia (e). Boxplots show quartiles and medians calculated from each image stack. Data points represent case means (females = circles, males = triangles). Stats = LMEM (Variable ∼ Plaque_vs_no-plaque*Group + 1|Case/ROI_number). *p*-values were calculated from post-hoc testing with Tukey correction for multiple comparisons.

**Figure 3:**
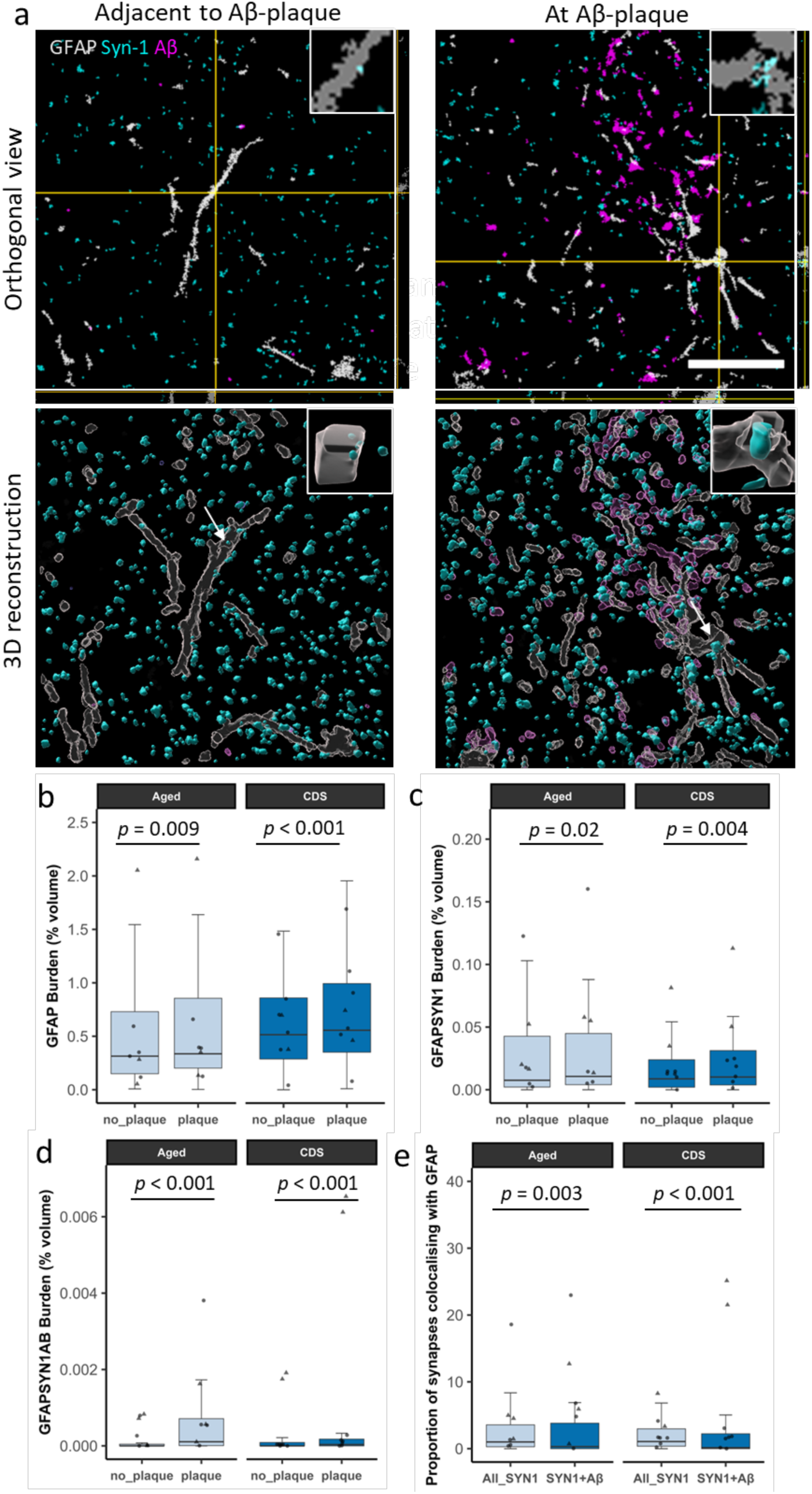
Amyloid-beta pathology induces increased synaptic engulfment by astrocytes in the feline brain 3. **a)** Segmented confocal images of CDS-affected feline parietal cortex, showing a region with amyloid-beta pathology and a paired image acquired approximately one field of view away in the same cortical layer in an area without pathology. Tissue was immunostained for synapses (synapsin-1, cyan), amyloid-beta (magenta), and astrocytes (GFAP, grey). Scale bar = 20 µm. Inserts = 5 µm * 5 µm. The top row of images shows orthogonal views. The bottom row of images shows 3D reconstructions generated using IMARIS and have been rotated along the z-axis, to further demonstrate that synaptic protein is within astrocytes. Inserts show close up of colocalisation highlighted by arrow. **b-e)** Colocalisation analysis reveals that the image stacks acquired in regions with amyloid-beta pathology show increased astrogliosis (GFAP burden) (b), colocalisation between astrocytes and synapses (GFAP and Synapsin-1 burden) (c), and triple colocalisation between astrocytes, synapses and amyloid-beta (GFAP, synapsin-1 and amyloid-beta burden) (d). Finally, when analysing all image stacks, synapses colocalising with amyloid-beta are less likely to be engulfed by astrocytes (e). Boxplots show quartiles and medians calculated from each image stack. Data points represent case means (females = circles, males = triangles). Stats = LMEM (Variable ∼ Plaque_vs_no-plaque*Group + 1|Case/ROI_number). *p*-values were calculated from post-hoc testing with Tukey correction for multiple comparisons.

### Amyloid-beta pathology induces increased synaptic engulfment by microglia in the feline brain

To investigate if amyloid-beta pathology may contribute to synaptic loss via the induction of aberrant synaptic pruning by microglia in the feline brain, an ROI was selected where there was amyloid-beta pathology and then selected a paired ROI approximately one field of view over in an area without amyloid-beta pathology but still within the same cortical layer (**Figure 2a)**. LMEM of the data (Variable ∼ Plaque_vs_no-plaque*Group + 1|Case/ROI_number) found that in regions of amyloid-beta pathology there was microgliosis (increased IBA1 burden) in both the aged (*t*(122) = 2.798, *p* = 0.006) and CDS group (*t*(122) = 5.089, *p* < 0.001)(**Figure 2b)**. In the aged (*t*(122) = 4.014, *p* < 0.001) and CDS group (*t*(122) = 4.780, *p* < 0.001) there was also increased internalisation of synapses by microglia (synapsin-1 and IBA1 colocalisation) around amyloid-beta plaques (**Figure 2c**). Further, near amyloid-beta plaques we found increased internalisation of synapses colocalising with amyloid-beta by microglia (synapsin-1, 4G8, and IBA1 triple colocalisation) in both the aged (*t*(122) = 8.353, *p* < 0.001) and CDS group (*t*(122) = 8.678, *p* < 0.001)(**Figure 2d)** – suggesting that microglia may be engulfing synapses that contain amyloid-beta. When analysing all image stacks, including those taken at an amyloid plaque and away from an amyloid plaque, we found that synapses colocalising with amyloid-beta were more likely to colocalise with microglia in both the aged (*t*(243) = 5.974, *p* < 0.001) and CDS groups (*t*(247) =15.464, *p* < 0.001)(**Figure 2e)**. Finally, we compared the IBA1 burden, IBA1 and synapsin-1 colocalisation, IBA1 and 4G8 colocalisation, and IBA1, 4G8 and synapsin-1 colocalisation between the young, aged and CDS groups. In this analysis, the 10 image stacks with the highest levels of amyloid-beta pathology were selected for each case. For all readouts, we found an increase in the CDS group compared to the young group. We found no differences between the aged group and the young group or the aged group and the CDS group (**supplementary figure 1**).

### Amyloid-beta pathology induces increased synaptic engulfment by astrocytes in the feline brain

We observed astrogliosis, by way of increased GFAP burden, in areas with amyloid-beta plaques when compared to regions without plaques in both the aged (*t*(122) = 2.666, *p* < 0.009) and CDS groups (*t*(242) = 4.644, *p* < 0.001)(**Figure 3b**). In the aged (*t*(122) = 2.360, *p* = 0.02) and CDS groups (*t*(122) = 2.907, *p* = 0.004) there was also increased colocalisation of synapses (synapsin-1) with astrocytes (GFAP) around amyloid-beta plaques (**Figure 3c**). Additionally, there was increased triple colocalisation between astrocytes (GFAP), synapses (synapsin-1) and amyloid-beta (4G8) around amyloid-beta pathology in both the aged (*t*(122) = 6.969, *p* < 0.001) and CDS groups (*t*(122) = 5.193, *p* < 0.001)(**Figure 3d**) – again suggesting that astrocytes might be eliminating amyloid-beta containing synapses. However, in contrast to what we found with microglia, synapses colocalising with amyloid-beta were less likely to colocalise with astrocytes in both the aged (*t*(242) = 2.993, *p* = 0.003) and CDS groups (*t*(247) =4.079, *p* < 0.001)(**Figure 3e**). Finally, we compared the GFAP burden, GFAP and synapsin-1 colocalisation, GFAP and 4G8 colocalisation, and GFAP, 4G8 and synapsin-1 colocalisation from all of the image stacks between the young, aged and CDS groups (**supplementary figure 2**). In this analysis, the 10 image stacks with the highest levels of amyloid-beta pathology were selected for each case. When comparing between the CDS and young group, the CDS group had an increase in the burden of GFAP-positive astrocytes (*t*(22) =2.610, *p* = 0.0408)(**supplementary figure 2a**), GFAP and amyloid-beta colocalisation (*t*(22) =3.610, *p* = 0.0043)(**supplementary figure 2d**), and a trend for an increase in triple colocalisation between GFAP-positive astrocytes, amyloid-beta and synapsin-1 (*t*(22) =2.459, *p* = 0.0558)(**supplementary figure 2c**). No other comparisons showed no difference between groups.

## Discussion

In this study, we observe that amyloid-beta colocalises with synapses in the aged and CDS-affected feline brain and is associated with regional gliosis, as well as increased synaptic engulfment by both microglia and astrocytes.

Our finding that amyloid-beta colocalises with synapses in feline CDS is in line with previous findings in human AD brain and rodent models of amyloid pathology (Koffie *et al*., 2009, 2012; Jackson *et al*., 2019; Pickett *et al*., 2019; King *et al*., 2023) and supports the hypothesis that amyloid-beta accumulation contributes to behavioural and cognitive changes observed in feline CDS. The exact mechanisms by which amyloid-beta induces synaptotoxicity remain a matter of debate. Data from preclinical studies and human postmortem tissue have implicated the interaction of Aβ with several synaptic proteins in mediating its synaptotoxic effects, such as cellular prion protein (PrPC), transmembrane protein 97, metabotropic glutamate receptor 5 (mGluR5), and NMDA receptors (Laurén *et al*., 2009; Larson *et al*., 2012; Folch *et al*., 2018; Colom-Cadena *et al*., 2024).

Additionally, our finding that gliosis and increased synaptic engulfment by microglia and astrocytes occur in the vicinity of amyloid-beta plaques further supports the notion that amyloid-beta exerts a pathological effect within the feline brain. From our data, we are unable to determine whether glia are removing degenerating or functional synapses. In support of the latter, data from model systems of AD suggest that increased synaptic engulfment by microglia leads to synapse loss and cognitive dysfunction. Importantly, blocking synaptic engulfment in these models rescues cognitive function (Hong *et al*., 2016b; Shi *et al*., 2017; Bie *et al*., 2019). This indicates that microglia are engulfing functional synapses and thereby contributing to clinical progression. Furthermore, increased synaptic engulfment by microglia does not appear to be a common feature of neurodegenerative diseases, but rather a direct consequence of amyloid-beta accumulation. We recently demonstrated increased synaptic engulfment by both astrocytes and microglia in AD (Tzioras, Daniels, *et al*., 2023). However, in the brains of individuals who died with the primary tauopathies progressive supranuclear palsy (PSP) (McGeachan *et al*., 2024) and frontotemporal dementia caused by the MAPT 10+16 mutation (Dando *et al*., 2024), we did not observe increased microglial engulfment, and increased astrocytic engulfment was only observed in PSP. Our data showing that synaptic engulfment is greatest in the vicinity of amyloid-beta plaques, and the finding that amyloid-beta-containing synapses are more likely to be engulfed when compared to all synapses in the feline brain, further support this hypothesis. Therefore, feline CDS serves as a naturally occurring model of human AD, providing a valuable avenue for studying disease mechanisms and testing therapeutic interventions targeting amyloid-beta-induced synaptic loss. Studying feline CDS has the potential to enhance our understanding and management of feline CDS while simultaneously contributing to the development of treatments for human dementia.

For several pathological readouts, including microgliosis, astrogliosis, and synaptic engulfment by microglia, our findings show an increase when comparing the CDS (cognitive dysfunction syndrome) group to the young group. However, no significant differences were observed between the CDS group and age-matched controls, nor between the aged and young groups. These results suggest that the pathological changes observed in CDS may arise from an interaction between aging processes and these mechanisms, rather than being solely attributable to either factor independently. While the data highlight a potential link between these pathologies and CDS, the lack of distinction between CDS and age-matched controls makes it difficult to conclusively attribute these changes as causal in the development of CDS. However, the significant increase in gliosis and synaptic engulfment around plaques in our study and the previously reported significant increase in plaques in CDS indicate that these phenotypes my contribute to cognitive dysfunction in aging cats. It is also plausible that increased synaptic accumulation of amyloid-beta, gliosis, and increased synaptic engulfment represent a combination of disease mechanisms that collectively contribute to the neurodegeneration and behavioural changes observed in CDS. However, these factors may not be solely sufficient to explain the development of CDS in aged cats. Additionally, one limitation of our study is the relatively small sample size. We used all available samples, but obtaining body donations from owners is a sensitive topic and presents a logistical challenge. It is possible that our study is underpowered to detect smaller effect sizes between the young and aged groups or the aged and CDS groups. Despite lack of significance between CDS and aged cats in the measurements performed in this study, there is a stepwise increase in pathological features we measured between young, aged, and CDS animals and significant main effects of ANOVAs in many measures, indicating that further research with a larger sample size is warranted to explore this question in more depth. The resolution of confocal microscopy is limited by the diffraction limit of light, making it difficult to confidently resolve individual synapses. Future studies should aim to confirm the presence of amyloid-beta within synapses and determine the proportion affected using higher-resolution techniques, such as immunogold or correlative light and electron microscopy. We did not have access to appropriately prepared samples for such analysis.

In conclusion, our data provide insight into mechanisms by which amyloid-beta pathology may lead to synaptic dysfunction and loss, revealing a potential link between age-related amyloid-beta deposition and the behavioural and cognitive changes observed in feline cognitive dysfunction syndrome.

## Acknowledgements

The confocal microscope used for these experiments was generously funded by Alzheimer’s Research UK (ARUK-EG2016A-6) and a Wellcome Trust Institutional Strategic Support Fund at the University of Edinburgh. J.L. was funded by UCB Biopharma, as was the Oxford Nanoimager. This work was supported by funding to Dr Robert McGeachan via the Wellcome Trust, as part of the Edinburgh Clinical Academic Track for Veterinary Surgeons (225442/Z/22/Z) and by the UK Dementia Research Institute [award number UKDRI-Edin005] through UK DRI Ltd, principally funded by the UK Medical Research Council.

## Data Accessibility Statement

Upon acceptance for publication, all spreadsheets and statistical analysis files will be shared on Edinburgh Datashare. Raw images and data available from the lead authors upon reasonable request.

## Supplementary data

**Supplementary figure 1:**
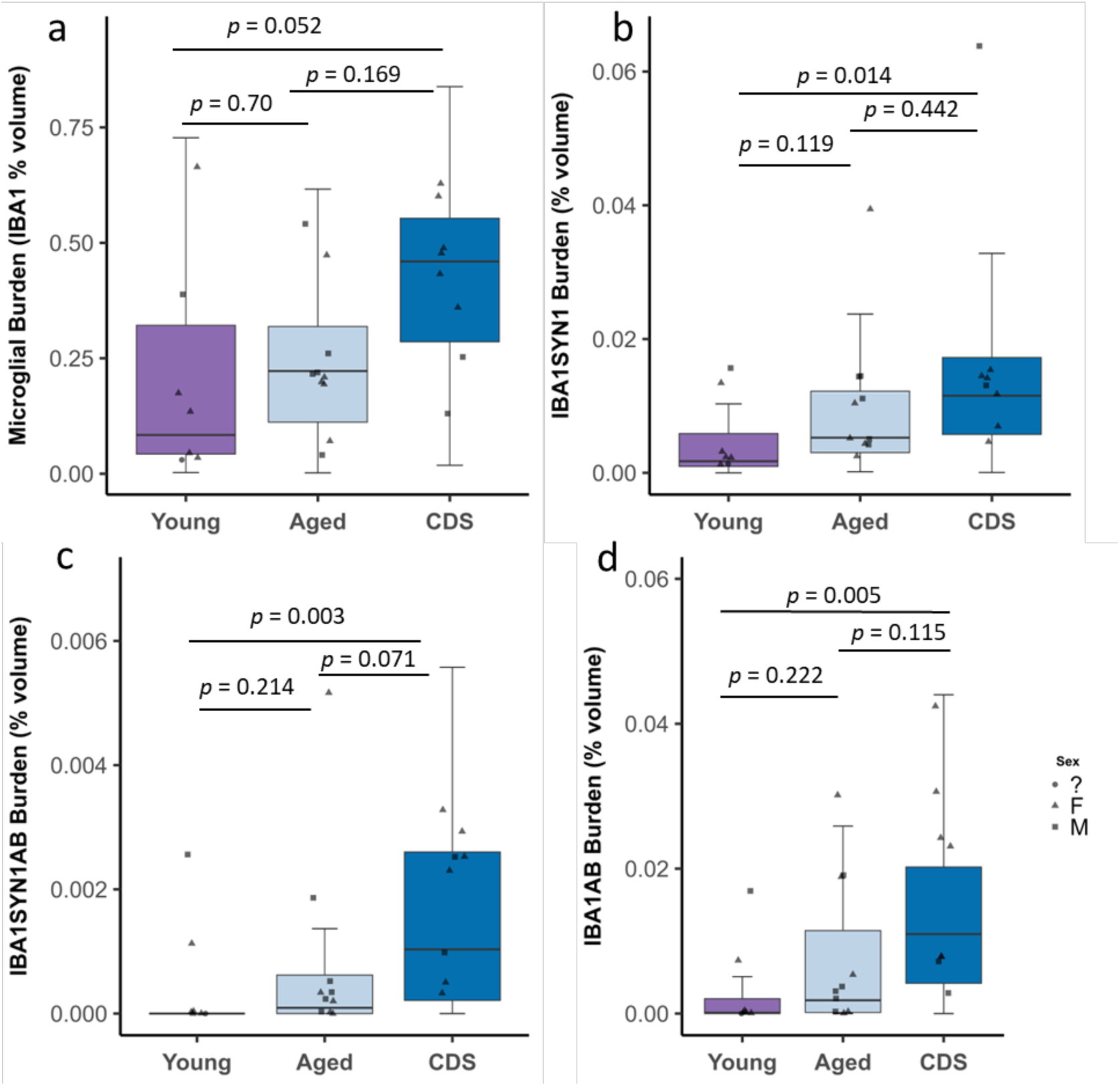
Effect of ageing and CDS on microglia – synapse interactisons in feline parietal cortex. **a)** Quantitative analysis reveals a trend for an increase in IBA1 burden in CDS group compared to the young (*t*(22) =2.494, *p* = 0.052) but no difference when compared to the aged group (*t*(22) = 1.877, *p* = 0.169). There was no difference between aged and young group (*t*(22) =0.812, *p* = 0.70) **b)** Quantitative analysis reveals an increase in IBA1 and synapsin-1 colocalisation in CDS group compared to the young (*t*(22) =3.113, *p* = 0.0135) but no difference when compared to the aged group (*t*(22) = 1.243, *p* = 0.4416). There was no difference between aged and young group (*t*(22) =2.074, *p* = 0.1186) **c)** Quantitative analysis reveals an increase in IBA1, synapsin-1 and amyloid-beta triple colocalisation in CDS group compared to the young (*t*(22) =3.796, *p* = 0.0027) and a trend towards an increase when compared to the aged group (*t*(22) = 2.337, *p* = 0.0714). There was no difference between the aged and young groups (*t*(22) =1.737, *p* = 0.2143) **d)** Quantitative analysis reveals an increase in IBA1 and amyloid-beta colocalisation in CDS group compared to the young (*t*(22) =3.548, *p* = 0.0049 but no difference when compared to the aged group (*t*(22) = 2.089, *p* = 0.1153). There was no difference between aged and young group (*t*(22) =1.716, *p* = 0.2219 **a-d)** Boxplots show quartiles and medians calculated from each image stack. Data points represent case means (sex unknown = circles, females = triangles, males = squares). Statistics: LMEM (variable ∼ Group + 1 | case). *p*-values were calculated from post-hoc testing with Tukey correction for multiple comparisons.

**Supplementary figure 2:**
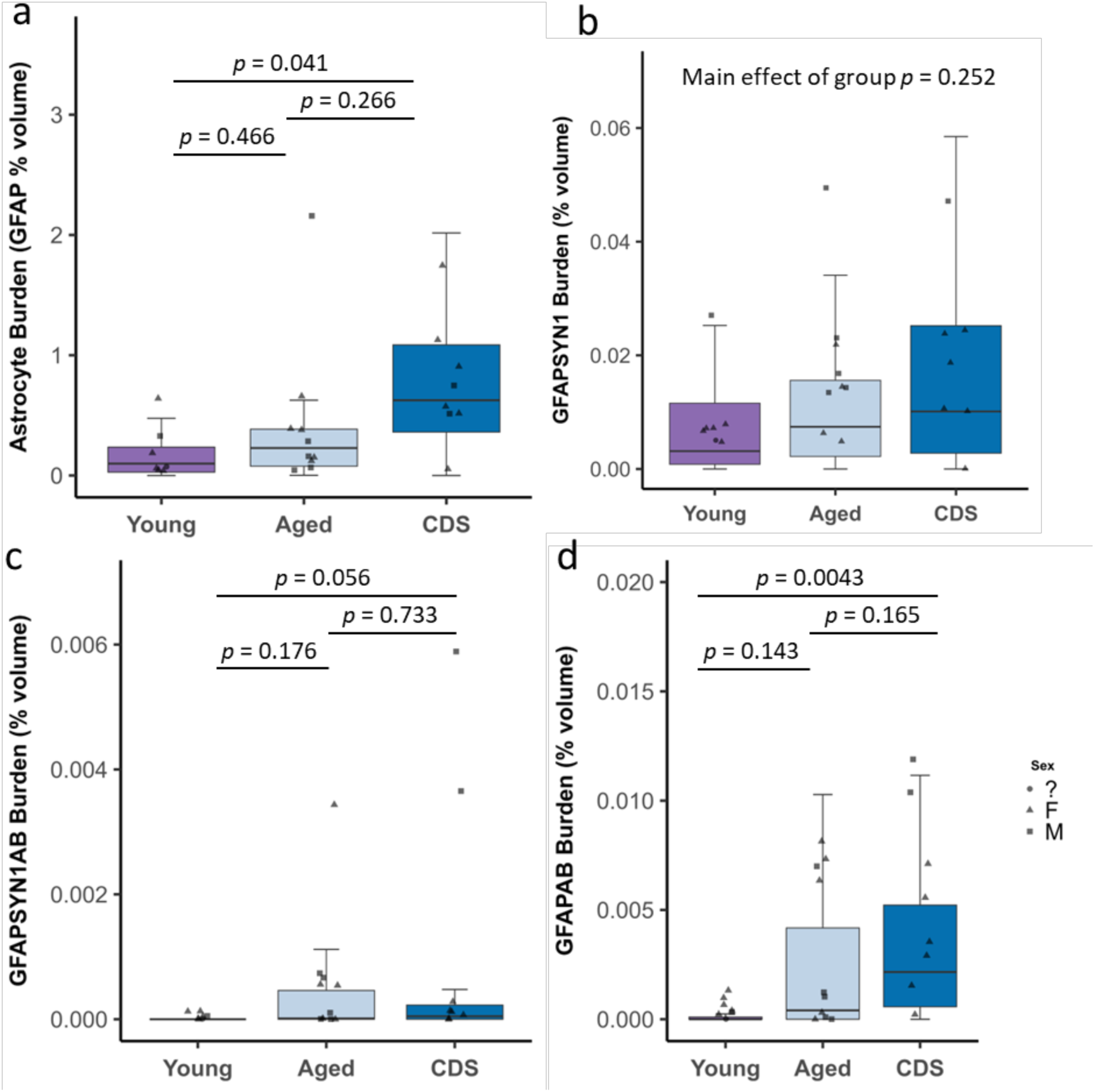
Effect of ageing and CDS on astrocyte – synapse interactions in the feline parietal cortex. **a)** Quantitative analysis reveals an increase in GFAP burden in CDS group compared to the young (*t*(22) =2.610, *p* = 0.0408) but no difference when compared to the aged group (*t*(22) = 1.602, *p* = 0.266). There was no difference between aged and young group (*t*(22) =1.199, *p* = 0.4661) **b)** Anova aner linear mixed effect modelling of the data reveals no main effect of group on the colocalisation between astrocytes and synapsin-1 (F[2/21.96] = 1.4674, *p* = 0.2523). As there was no main effect, pairwise comparisons were not performed. **c)** Quantitative analysis reveals a trend for an increase in GFAP, synapsin-1 and amyloid-beta triple colocalisation in CDS group compared to the young (*t*(22) =2.459, *p* = 0.0558) 0.0049 but no difference when compared to the aged group (*t*(22) = 0.756, *p* = 0.733). There was no difference between the aged and young groups (*t*(22) =1.855, *p* = 0.1757) **d)** Quantitative analysis reveals an increase in GFAP and amyloid-beta colocalisation in CDS group compared to the young (*t*(22) =3.610, *p* = 0.0043) but no difference when compared to the aged group (*t*(22) = 1.891, *p* = 0.1651). There was no difference between aged and young group (*t*(22) =1.972, *p* = 0.1428) a-d) Boxplots show quartiles and medians calculated from each image stack. Data points represent case means (sex unknown = circles, females = triangles, males = squares). Statistics: LMEM (variable ∼ Group + 1 | case). *p*-values were calculated from post-hoc testing with Tukey correction for multiple comparisons.

